# Oral Administration of Boldine Reduces Spare Nerve Injury-Induced Neuropathic Pain

**DOI:** 10.1101/2024.11.05.622128

**Authors:** Jiangping Pan, Carlos A. Toro, Christine Chow, Yorley Duarte, Juan C. Sáez, Christopher P. Cardozo, Wei Zhao

**Affiliations:** Spinal Cord Damage Research Center, James J. Peters Veteran Affairs Medical Center, Bronx, NY, United States; Department of Medicine, Icahn School of Medicine at Mount Sinai, New York, NY, United States; Bronx Veterans Medical Research Foundation, Bronx, NY, United States; Center for Bioinformatics and Integrative Biology, Facultad de Ciencias de la Vida, Universidad Andrés Bello, Santiago, Chile; Institute of Neuroscience, Centro Interdisciplinario de Neurociencia De Valparaíso, Universidad de Valparaíso, Valparaíso, Chile; Department of Rehabilitation and Human Performance, Icahn School of Medicine at Mount Sinai, New York, NY, United States

**Keywords:** neuropathic pain, nerve injury, nutraceuticals, microglia, inflammation

## Abstract

**Background:** Chronic pain is present in about 20% of the population and is a major burden to the health care system. About 30-40% of these patients report neuropathic pain. Neuropathic pain is defined as pain caused by injury or disease of the somatosensory nervous system. Current available treatments for neuropathic pain have limited efficacy and substantial side effects.

**Methods:** To address the need for more effective and safer treatments for neuropathic pain, this study aimed to test whether boldine, a naturally occurring alkaloid, could attenuate neuropathic pain in a murine model of spared nerve injury (SNI). Von Frey filament test, hot/cold plate test and dynamic weight bearing test were used to assess pain phenotypes following SNI.

**Results:** We found that boldine inhibited the lipopolysaccharide-induced overexpression of inflammatory markers in BV-2 microglial cells. Oral administration of boldine at 50 mg/kg body weight/day resulted in significant reduction of SNI-induced mechanical and thermal hypersensitivity. Boldine also corrected SNI-induced weight bearing deficits, which are an indication of spontaneous pain. Boldine significantly inhibited SNI-induced peripheral inflammation as indicated by reduced levels of inflammatory cytokines/chemokines in the serum. Immunofluorescence studies revealed that boldine reduced the number of reactive astrocytes and inhibited microglia activation in lumbar spinal cord.

**Conclusion:** Our findings suggest that boldine may be a promising therapeutic candidate for the treatment of neuropathic pain, possibly through inhibition of glia activation and neuroinflammation.

## Introduction

As described by the International Association for the Study of Pain, neuropathic pain is “pain caused by a lesion or a disease of the somatosensory nervous system”. It affects 7-10% of the general population globally [12]. Neuropathic pain can be central or peripheral, and tissue damage is not always apparent. Studies have shown that neuropathic pain has multiple adverse effects in both humans and rodents that include impaired sleep, depression, anxiety, and lower quality of life [4; 14; 36; 42]. Common causes of neuropathic pain include trigeminal neuralgia, post-amputation pain, persistent post-operative or post-traumatic pain, pain after peripheral nerve injury, painful diabetic neuropathy, cancer pain, and chemotherapy-induced peripheral neuropathy. Due to its subjective nature, diagnosing neuropathic pain is challenging. Typical characteristics of neuropathic pain include allodynia, where normally non-painful stimuli cause pain; hyperalgesia, an exaggerated response to painful stimuli; and spontaneous pain [2; 28; 34; 41].

Currently, the first line of choice for neuropathic pain treatment includes: anticonvulsants, tricyclic antidepressants, serotonin and norepinephrine re-uptake inhibitors, and topical agents [5]. Opioids, such as morphine and oxycodone, can be administered under careful medical supervision for very severe cases, although their long-term efficacy in treating neuropathic pain is uncertain, and they can be associated with severe side effects and risks, including sedation, respiratory depression, potential dependence and addiction [17]. Thus, there is an urgent need to develop novel, efficacious and safe interventions for treating neuropathic pain. Recently, the beneficial effect of some nutraceuticals and herbal remedies have gained attention in combating neuropathic pain [1]. Curcumin for instance, reduced neuropathic pain in streptozotocin-induced diabetic neuropathy [51] and peripheral nerve injury rodent models [25].

Boldine is the most abundant alkaloid isolated from the leaves and bark from boldo (*Peumus boldus*), a widely distributed evergreen tree native to Chile [48]. Boldo extracts containing boldine has been used since the last century in Europe and North/South America, for the treatment of digestive problems, headache, rheumatism and urinary tract inflammation [53].

Boldine has also been shown to have antioxidative activities that protect both biological and non-biological systems against free radical-mediated oxidative damage [45]. In addition, boldine exhibits hepatoprotective, anti-inflammatory and antipyretic activities in animal models [3; 8]. Recent studies conducted by us and others have shown the beneficial effect of boldine in neurological conditions associated with neuroinflammation including spinal cord injury [16; 56], delayed nerve repair [7], antiepileptic [43], and Alzheimer’s Disease [55], suggesting that boldine could pass the blood-brain barrier. Consistent with this interpretation, boldine accumulates in brain tissues after oral administration [30]. Considering the anti-inflammatory activity of boldine and the key role of neuroinflammation in the development and maintenance of neuropathic pain, we evaluated the potential of boldine to attenuate neuropathic pain in a mouse model of spared nerve injury.

## Material and Methods

### Reagents

Dulbecco’s Modified Eagle Medium (DMEM), fetal bovine serum (FBS), Dulbecco’s Phosphate-Buffered Saline (DPBS), penicillin and streptomycin, were purchased from GIBCO (Waltham, MA). Lipopolysaccharides (LPS), boldine, dimethyl sulfoxide (DMSO), triton X-100, and paraformaldehyde (PFA) were purchased from Sigma-Aldrich (St. Louis, MO). LPS was dissolved in PBS and boldine was first dissolved in DMSO then diluted in culture medium.

### Cell culture, treatment, cytotoxicity, and ELISA

BV-2 cells were obtained from AcceGen (Fairfield, NJ) and maintained in DMEM supplemented with 10% FBS and antibiotics (penicillin 100 U/mL, streptomycin 100 μg/mL) at 37°C with 5% CO_2_. Cells (in triplicates) were treated with 100 ng/mL LPS with or without boldine (25 or 50 µg/mL) for 6 hours, and 100 μL of the supernatant was sampled for cytotoxicity essay before cells were harvested for RNA extraction or ELISA. Cell viability was assessed using the CytoTox 96 Non-Radioactive Cytotoxicity Assay (Promega, Fitchburg, WI) following manufacturer’s instructions. For ELISA, BV-2 cells were harvested and lysed in RIPA buffer (Pierce Biotechnology, Waltham, MA) and protein concentration was measured using the BCA Protein Assay Kits (Pierce Biotechnology). IL-1β protein levels were assessed using the Mouse IL-1 beta/IL-1F2 Quantikine ELISA Kit (R&D Systems, Minneapolis, MN) following the manufacturer’s directions. Optical densities were measured using a SpectraMax iD5 plate reader (Molecular Devices, San Jose, CA).

### Animal, surgery, and boldine administration

Animals were maintained on a 12:12-h light/dark cycle in a temperature-controlled (20 ± 2°C) vivarium, with food and water *ad libitum*. All procedures were approved by the Institutional Animal Care and Use Committee at James J. Peters Veterans Affairs Medical Center (#1585874) and conformed to all guidelines and regulations for the protection of the welfare of animals. C57BL6 mice were purchased from the Jackson Laboratory (Bar Harbor, ME) and pair-housed. Spared nerve injury (SNI) surgery was conducted on the left hindlimb under anesthesia (3-5% isoflurane) by cutting the peroneal and tibial nerves and leaving an intact sural nerve. Sham animals underwent a similar procedure where the branches of the sciatic nerve were exposed but left intact. Boldine was administered to the animal right after surgery via a customized chow (Research Diets, New Brunswick, NJ) to reach a dose of 50 mg/Kg BW/day. A control diet (OpenStandard Diet With 15 Kcal% Fat, Research Diets) was provided to sham-operated or SNI group. Animals were euthanized at 15 days post-surgery. Lumbar spinal cords (L3-5) were dissected and stored at -80 °C for further analysis. Blood samples were collected and left at room temperature for 1 hour to allow it to clot. Blood was then centrifuged at 1,000 x g for 10 minutes and supernatant (serum) was stored at -80 °C for further analysis.

### RNA Isolation, cDNA synthesis and quantitative PCR

BV-2 cells were harvested and RNAs were extracted using the RNeasy Kit (Qiagen, Hilden, Germany) following the manufacturer’s directions. Frozen tissues were homogenized in TRI-Reagent (Sigma-Aldrich, St. Louis, MO) using a FastPrep-24™ 5G Bead Beating Grinder and Lysis System (MP Biomedicals, Solon, OH). Total RNA was isolated following the manufacturer’s instructions. RNA concentrations were measured using a NanoDrop Spectrophotometer (Thermo Fisher Scientific, Waltham, MA). One microgram of total RNA was reverse transcribed with High-Capacity cDNA Reverse Transcription Kit using random hexamers in a 20 μL reaction according to the manufacturer’s protocol. Quantitative PCR (qPCR) was performed using the Taqman Universal PCR Master Mix on a ViiA 7 Real-Time PCR System. Relative expression levels were calculated using the 2^−ΔΔCt^ method with hypoxanthine phosphoribosyltransferase 1 (HPRT) as an internal control. A list of the Taqman probes used in this study is provided in Supplemental Table 1 (all from Applied Biosystems, Waltham, MA).

### Behavioral Tests

Mechanical pain responses were assessed by Von Frey filament test performed on the ipsilateral and contralateral hind paws [15]. Briefly, The Von Frey hair (Bioseb, Vitrolles, France) is inserted perpendicularly from below an elevated wire platform into the plantar surface of the hind paw, starting with the smallest gauge filament. The pressure was increased slowly, until the hair just began to bend. The test was then repeated with the next largest gauge filament until withdrawal responses including paw flicking, paw lifting, paw licking or possibly vocalizing was observed. The withdrawal threshold, defined as the smallest gauge wire that produced a withdrawal response, was recorded.

Thermal pain responses were measured using a hot/cold pate apparatus (Maze Engineers, Skokie, IL). Mice were allowed to move freely on a hot/cold plate enclosure for a maximum of 30 seconds. For assessing pain responses to heat, plate temperature was set at 50 ^°^C. For testing pain responses to cold, plate temperature was set at 15 ^°^C. Once paw retraction, paw licking, or escaping behaviors were observed, mice were immediately taken out of the testing apparatus and the time latency was recorded [63].

Spontaneous pain responses were measured using the Dynamic Weight Bearing 2.0 system (DWB2.0) (Bioseb, Vitrolles, France) [50]. Briefly, the animal was placed into the DWB enclosure for a test duration of 5 minutes. The animal was recorded from above using a high-definition camera and the captured video feed was analyzed in real-time using tracking software. An observer manually validated each automatically presumed paw position to avoid subsequent error identification afterward.

### Multiplex ELISA

MILLIPLEX® Mouse Cytokine/Chemokine Magnetic Bead Panel (Millipore, Billerica, MA) was used to measure the levels of 32 cytokines/chemokines in mouse serum as previously described [59; 60; 66]. Briefly, 25 µL of serum sample was diluted with assay buffer and incubated with magnetic beads overnight at 4°C. The samples were then washed, incubated with detection antibodies for 1 hour at room temperature followed by streptavidin-phycoerythrin incubation for 30 minutes. The plate was run on a Luminex 200 instrument (Millipore, Billerica, MA) and the median fluorescent intensity was used to calculate the concentration of each cytokine/chemokine.

### Immunostaining and microscopy

At 15 dpi, mice were transcardially perfused with saline followed by 4% paraformaldehyde (PFA). Lumbar spinal cords (L3-5) were harvested and transversely sectioned using a LEICA CM3050 Cryostat (LEICA, Wetzlar, Germany). For immunostaining, slides were first incubated with blocking buffer (1 x PBS + 5% goat serum + 0.3% Triton X-100) for 1h at room temperature, followed by incubation with primary antibodies of astrocyte marker glial fibrillary acidic protein (GFAP, Abcam, Cambridge, UK) or microglia marker IBA1 (Abcam) overnight at 4°C. After incubation, the slides were washed 3 times in PBS and incubated with goat anti-mouse Alexa Fluor 488 secondary antibody (Abcam) for 1 hour at room temperature. Slides were then washed with PBS three times and mounted with VECTASHIELD Antifade Mounting Medium (Vector Laboratories, Newark, CA). Images were acquired using the ZEISS LSM 700 confocal imaging system (Carl Zeiss, Oberkochen, Germany) with 20× objective. Mean fluorescent intensity was quantified using ImageJ.

### Statistical analysis

Data were analyzed using GraphPad Prism 10 (San Diego, CA) and were presented as the mean ± standard error of the mean (SEM). Two-way repeated-measures analysis of variance (ANOVA) followed by multiple comparison test were performed to compare the mean differences in mechanical thresholds, thermal thresholds and weight bearing among the experimental groups. One-way ANOVA followed by multiple comparison test was performed for the comparison of mean difference in cytotoxicity and RNA or protein expression levels among experimental groups. p-value of < 0.05 was considered statistically significant.

## Results

### Boldine inhibits LPS-induced inflammation in microglial cells

Neuroinflammation acts as both a driver and amplifier of pain states, particularly in chronic conditions. Boldine has been shown to exhibit anti-inflammatory activities [3; 8]. To test whether boldine could reduce neuroinflammation, BV-2 microglia cells were treated with LPS with or without boldine. After treatment, cell medium was collected and subjected to a cell viability assay. Neither the 25 µg/mL or the 50 µg/mL dose of boldine showed any cytotoxicity (Supplemental Figure 1A). Cells were then harvested and subjected to gene expression analysis by quantitative PCR. LPS treatment led to significantly elevated mRNA levels of chemokines (CCL2, CCL3 and CCL5), cytokines (IL-1β, IL-6, and IL-18), macrophage and microglia marker CD11b, toll-like receptors (TLR2 and TLR3), and NOD-, LRR- and pyrin domain-containing protein 3 (NLRP3) (Figure 1). Co-incubation with boldine inhibited the LPS-induced upregulation of the above-mentioned molecules, suggesting a beneficial role of boldine in LPS-induced inflammation. We also found that IL-1β protein levels were drastically increased after LPS treatment, which was prevented by boldine co-treatment (Supplemental Figure 1B).

**Figure 1.**
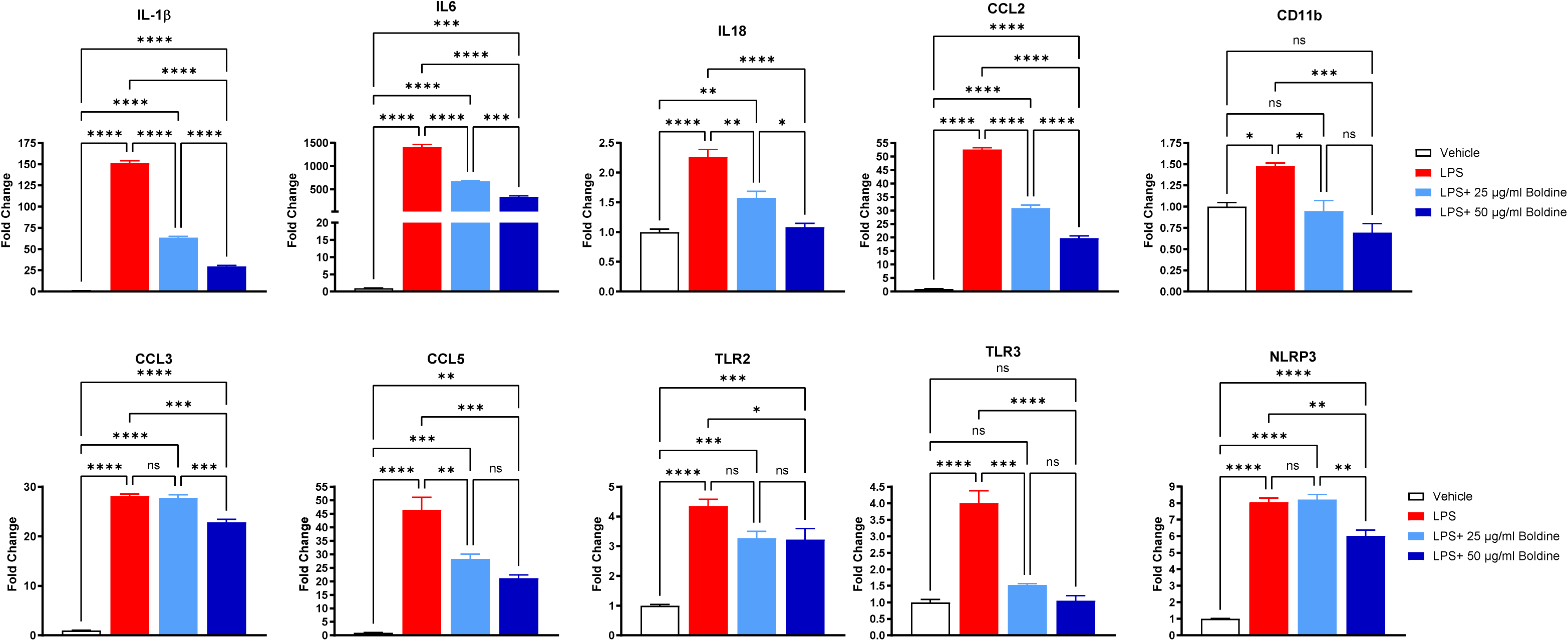
Boldine prevents LPS-induced inflammation *in vitro*. BV-2 microglia cells were treated with 100 ng/mL lipopolysaccharides (LPS) with or without boldine (25 µg/mL or 50 µg/mL) for 6 hours. Cells were harvested and subjected to RNA extraction followed by cDNA synthesis and quantitative PCR using standard Taqman methods. Taqman probes include CCL2, CCL3, CCL5, CD11b, IL-1β, IL-6, IL-18, TLR2, TLR3, and NLRP3. Relative levels were calculated using the 2^−ΔΔCt^ method with HPRT as an internal control. Data represent Mean ± SEM, N = 3 per condition, * p < 0.05, ** p < 0.01, *** p < 0.001, **** p < 0.0001 by one-way ANOVA followed by Tukey post-tests.

### Boldine alleviates SNI-induced mechanical allodynia

To assess the role of boldine in peripheral nerve injury-induced neuropathic pain, we conducted studies in a mouse model of SNI, which produces persistent peripheral hypersensitivity to mechanical stimuli [13]. Briefly, 10 weeks old female C57BL6 mice were subjected to SNI or sham surgery. Customized diet containing boldine or control diet was administered immediately after surgery. Pain phenotyping tests including Von Frey filament test, hot/cold plate test and dynamic weight bearing test were performed before injury for baseline and starting from 3 days after injury (Figure 2A, experiment design). The dosage of boldine was monitored by weekly measurement of food consumption (Figure 2B). The average food intake is 2.5 ∼ 2.7 gram per animal per day, which is equivalent to 50 ∼ 54 mg/kg BW/day of boldine. No significant difference in food intake was observed among experimental groups (Figure 2B).

**Figure 2.**
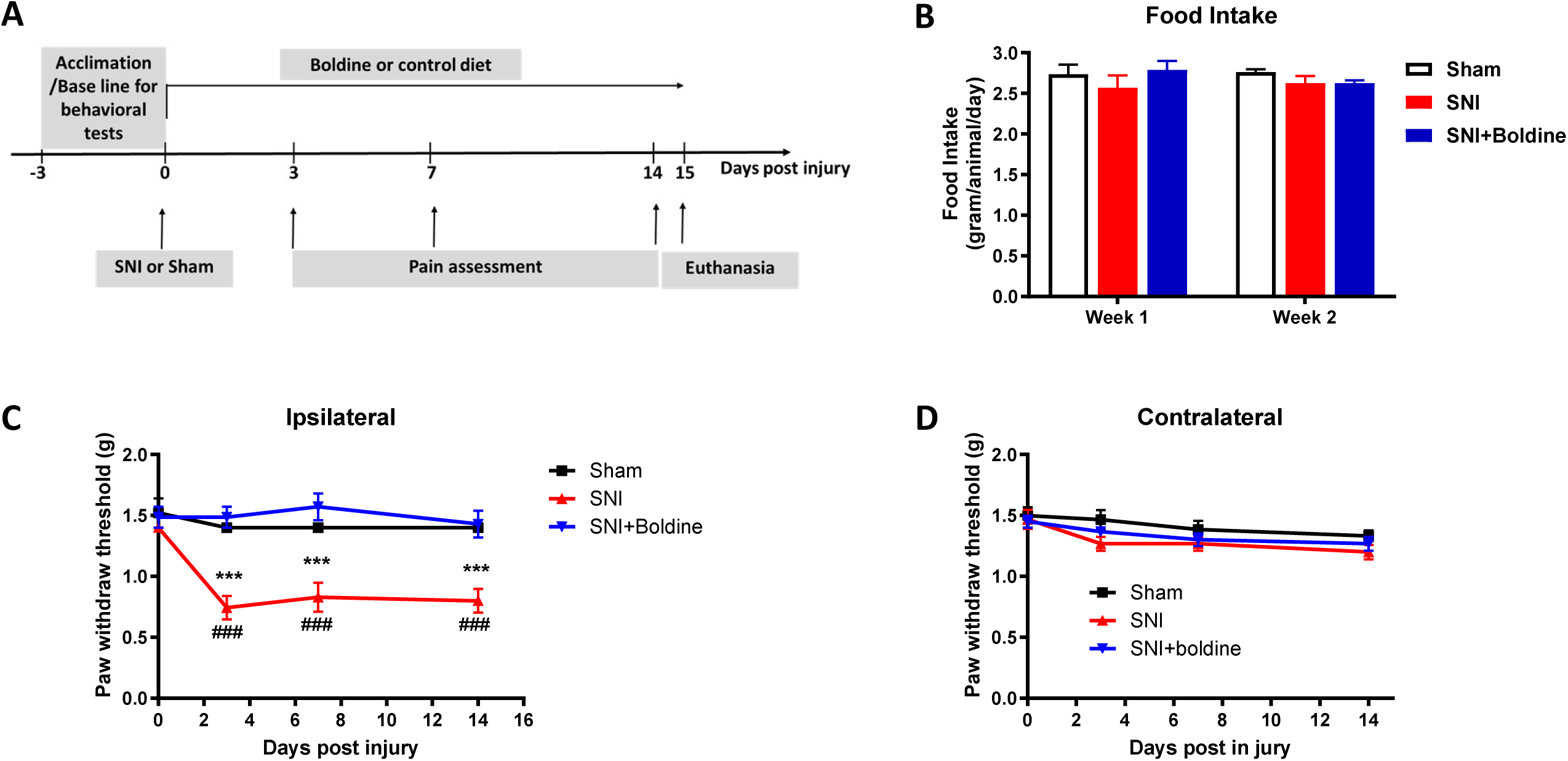
Boldine prevents spared nerve injury-induced mechanical allodynia. Female C57BL6 mice were subjected to spared nerve injury (SNI). Boldine was administered right after injury via a customized diet (Research Diet) to reach a dose of 50 mg/kg/day. A control diet was provided to sham or SNI group. Von Frey filament tests were performed to evaluate mechanical pain responses. (A) Diagram of experiment design; (B) Food intake was monitored weekly and average daily intake was calculated and plotted in GraphPad Prism 10. (C) Withdraw threshold to mechanical stimuli on the ipsilateral hind paw before and post injury; (D) Paw withdraw threshold to mechanical stimuli on the contralateral hind paw. Data represent mean ± SEM, N = 8-10 per group, *** p < 0.001 (SNI vs. sham), ### p < 0.001 (SNI+boldine vs. SNI) by two-way ANOVA with Tukey post-tests

As expected, SNI significantly induced mechanical pain hypersensitivity on the ipsilateral hindlimb at 3-, 7-, and 14-days post injury (dpi) when compared to sham-operated animals. Oral administration of boldine completely prevented this injury-induced mechanical allodynia (Figure 2C). There was no change of responses to mechanical stimuli after nerve injury or boldine treatment on the contralateral hind limb (Figure 2D), suggesting that the boldine-mediated prevention of mechanical-induced allodynia is specific to the region innervated by the injured nerve. We also observed a similar effect of boldine in preventing SNI-induced mechanical hypersensitivity in male mice (Supplemental Figure 2). We choose to focus on female mice for subsequent studies because women are more likely to have chronic pain and high-impact chronic pain compared with men [65].

### Boldine reduces SNI-induced thermal pain

Rodents with SNI often exhibit increased sensitivity to thermal stimuli, manifesting as decreased withdrawal latencies in response to heat or cold compared to control animals [13]. To test the effect of boldine in altering thermal pain phenotypes, we performed the hot plate test at and the cold plate test following SNI. At 7- and 14-dpi, SNI animals developed increased sensitivities to both warm (Figure 3A) and cold stimuli (Figure 3B). Administration of boldine significantly attenuated the SNI-induced hypersensitivity to warm and cold temperatures.

**Figure 3.**
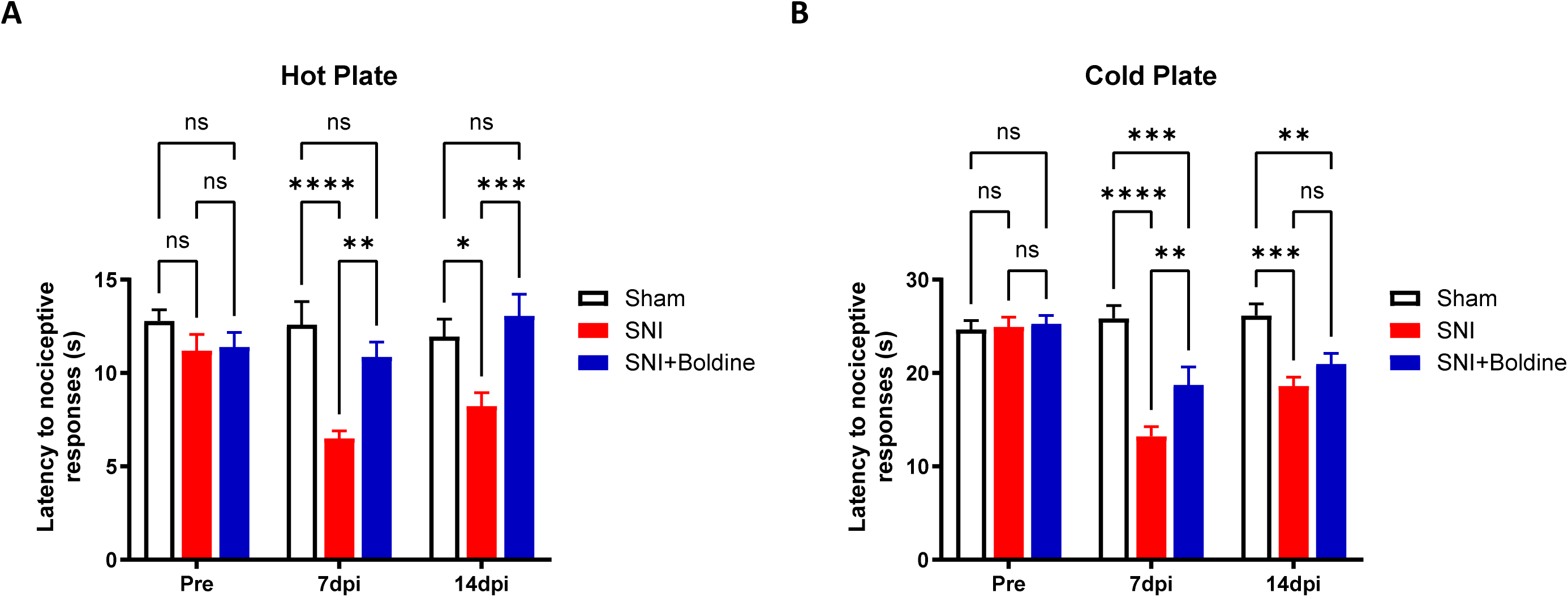
Boldine reduces SNI-induced thermal pain hypersensitivity. Hot/cold plate tests were performed to evaluate thermal pain responses before (Pre) and after (7dpi and 14dpi) injury in sham animals (Sham) and animals with (SNI + Boldine) or without (SNI) boldine treatment. (A) Latency to nociceptive responses by hot plate test (50°C). (B) Latency to nociceptive responses by cold plate test (15°C). Data represent Mean ± SEM, N = 8-10 per group, * p < 0.05, ** p < 0.01, *** p < 0.001 by Two-way ANOVA with Tukey post-test.

### Boldine prevents SNI-induced weight bearing deficits

Alterations in weight-bearing behavior can be indicative of pain, including those resulting from peripheral nerve injury. In the context of SNI, changes in weight-bearing distribution are often observed as part of the pain-related behavioral phenotype [44]. To evaluate the effect of boldine on potential SNI-induced weight bearing deficits, we performed weight bearing tests using the dynamic weight bearing 2.0 system. As compared to sham-operated mice, injured mice tended to use the ipsilateral hindlimb less for weight support at 3 and 7 dpi, and boldine administration significantly alleviated the SNI-induced weight bearing deficit at 7dpi (Figure 4A). We also measured some additional parameters including the hindlimb weight support area ratio and rearing duration. Rearing behavior, where a mouse stands on its hind legs with its forelimbs raising and head extending, has been used as a spontaneous pain indicator in mice [62]. SNI led to significantly decreased ipsilateral/contralateral weight support area ratio (Figure 4B) and markedly lower rearing duration (Figure 4C). However, the administration of boldine did not prevent SNI-induced deficits in weight support area or rearing duration (Figure 4B and C).

**Figure 4.**
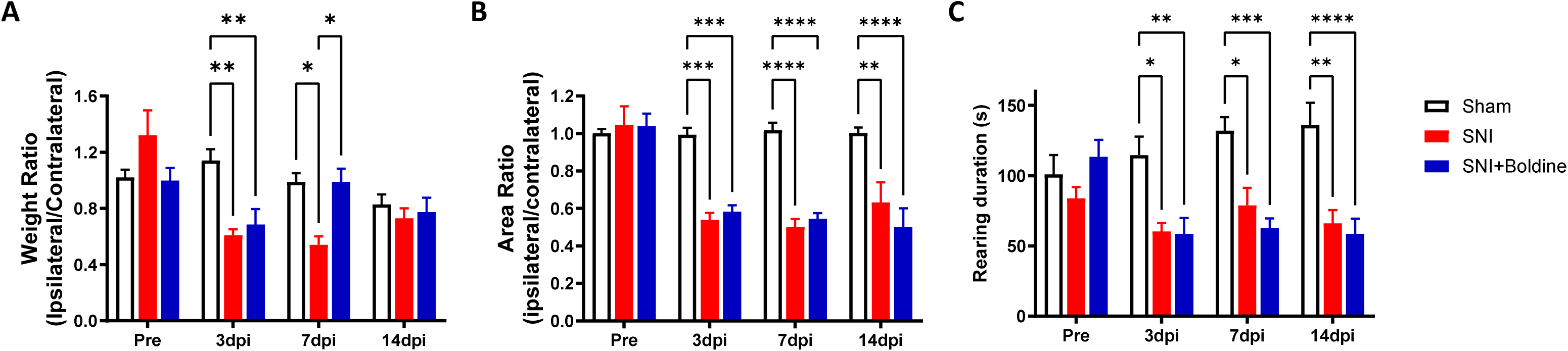
Effect of boldine on SNI-induced weight bearing deficits. Assessment of spontaneous pain measuring the changes in postural equilibrium was achieved using the Dynamic Weight Bearing System 2.0 (Bioseb) in sham animals (Sham) and animals with (SNI + Boldine) or without (SNI) boldine treatment. (A) Weight support ratio of the ipsilateral hind limb vs. the contralateral hind limb; (B) Support area ratio of the ipsilateral hind limb vs. the contralateral hind limb; (C) Rearing duration. Data represent Mean ± SEM, N = 8-10 per group, *p < 0.05, ** p < 0.01, *** p < 0.001, **** p < 0.0001 by two-way ANOVA with Tukey post-tests.

### Boldine reduces SNI-induced peripheral inflammation

To evaluate the effect of boldine on peripheral inflammation, serum samples were subjected to a multiplex ELISA using the MILLIPLEX® Mouse Cytokine/Chemokine Magnetic Bead Panel (Millipore). SNI induced the elevation of interleukin-12 (IL-12p40), interleukin-15 (IL-15), macrophage colony-stimulating factor (M-CSF) and chemokine ligand 5 (CCL5) in the serum, which was prevented by boldine treatment (Figure 5).

**Figure 5.**
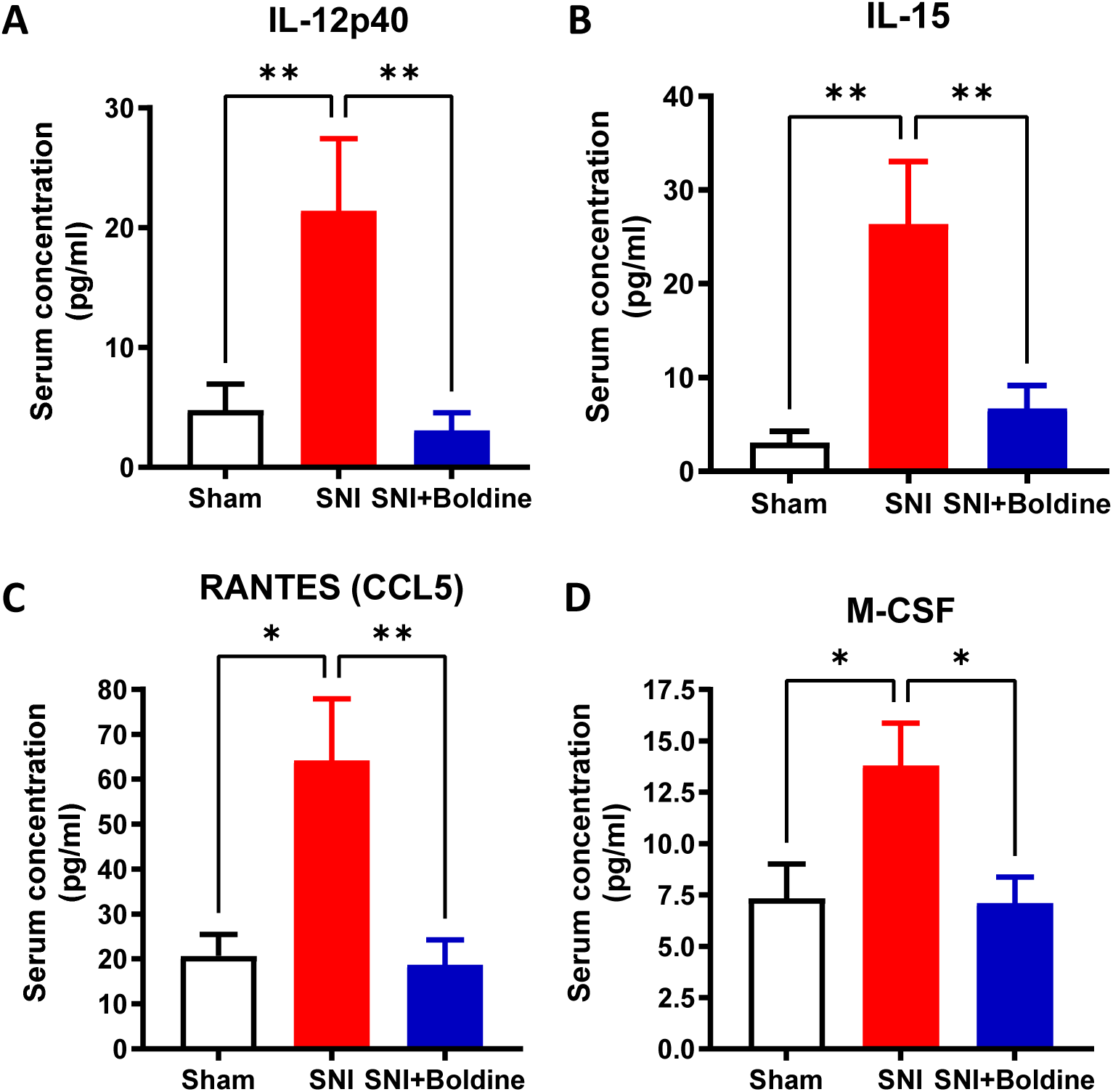
Peripheral inflammation changes after SNI and boldine treatment. Serum samples from sham (Sham) animals and mice subjected to spared nerve injury (SNI) or SNI plus 50 mg/kg boldine (SNI+Boldine) collected at 15 dpi were subjected to a multiplex ELISA using the MILLIPLEX® Mouse Cytokine/Chemokine Magnetic Bead Panel (Millipore) and the median fluorescent intensity was used to calculate the concentration of each cytokine/chemokine. (A) IL-12p40; (B) IL-15; (C) CCL5; (D) M-CSF. Data represent Mean ± SEM, N = 8-10 per group, * p < 0.05, ** p < 0.01 by one-way ANOVA with Tukey post-test.

### Boldine inhibits glia activation in the lumbar spinal cord

In pain conditions, nociceptive signals are transmitted from peripheral nerves to the spinal cord. Glial cells in the spinal cord dorsal horn play complex and dynamic roles in pain processing. Both activated microglia and reactive astrocytes have been implicated in pathogenesis of pain following nerve injuries [9; 10; 24; 31; 39]. To test the effect of boldine on glia activation, transverse lumbar spinal cord sections were immuno-stained with the astrocyte marker GFAP or the microglia marker IBA1. As compared to sham-operated mice, significantly more reactive astrocytes were detected in the lumbar spinal cord after SNI (Figure 6A), and boldine treatment prevented the increase in number of GFAP-positive cells in the ipsilateral dorsal horn (Figure 6B) and ventral horn (Figure 6C). Consistent with previous reports [20; 46], microglia in the spinal cord dorsal horn and ventral horns were activated after SNI (Figure 7A), and boldine treatment significantly reduced the SNI-induced microglia proliferation/activation in the ipsilateral dorsal horn (Figure 7B) and ventral horn (Figure 7C).

**Figure 6.**
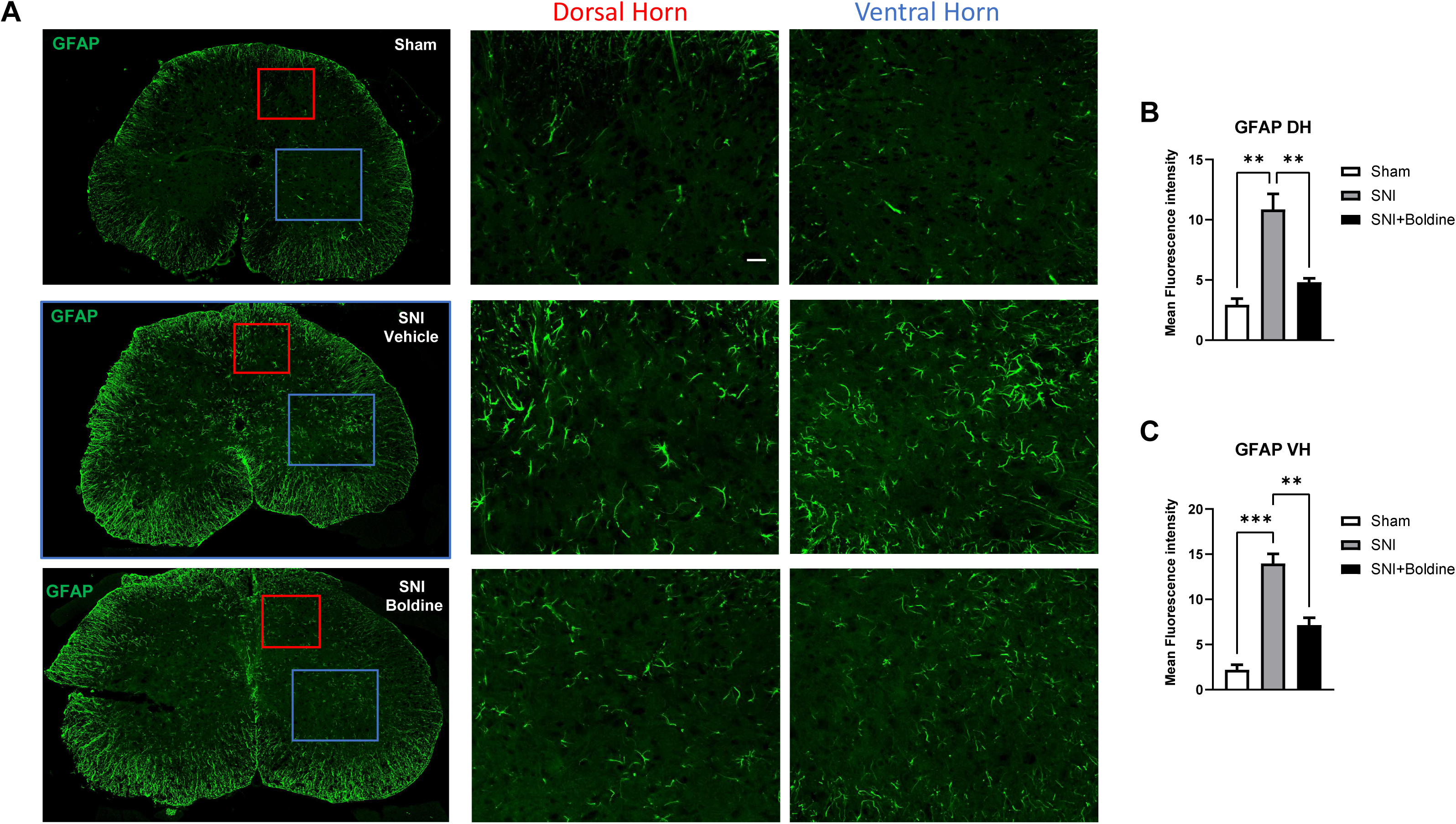
Boldine decreases levels of reactive astrocytes in the lumbar spinal cord after SNI. Sixteen micrometer thick transverse lumbar spinal cord sections were immuno-stained with astrocyte marker GFAP to evaluate the effect of SNI and boldine treatment on glia activation. Images were acquired using the ZEISS LSM 700 confocal imaging system with 20 × objective. (A) Representative images of the lumbar spinal cord. Red box: ipsilateral dorsal horn; Blue box: ipsilateral ventral horn. Scale bar: 10 µm. Quantification of GFAP expression at the (B) ipsilateral dorsal horn (DH) and (C) ipsilateral ventral horn (VH). Data represent Mean ± SEM, N = 3 mice per group. ** p < 0.01, *** p < 0.001 by one-way ANOVA with Tukey post-test.

**Figure 7.**
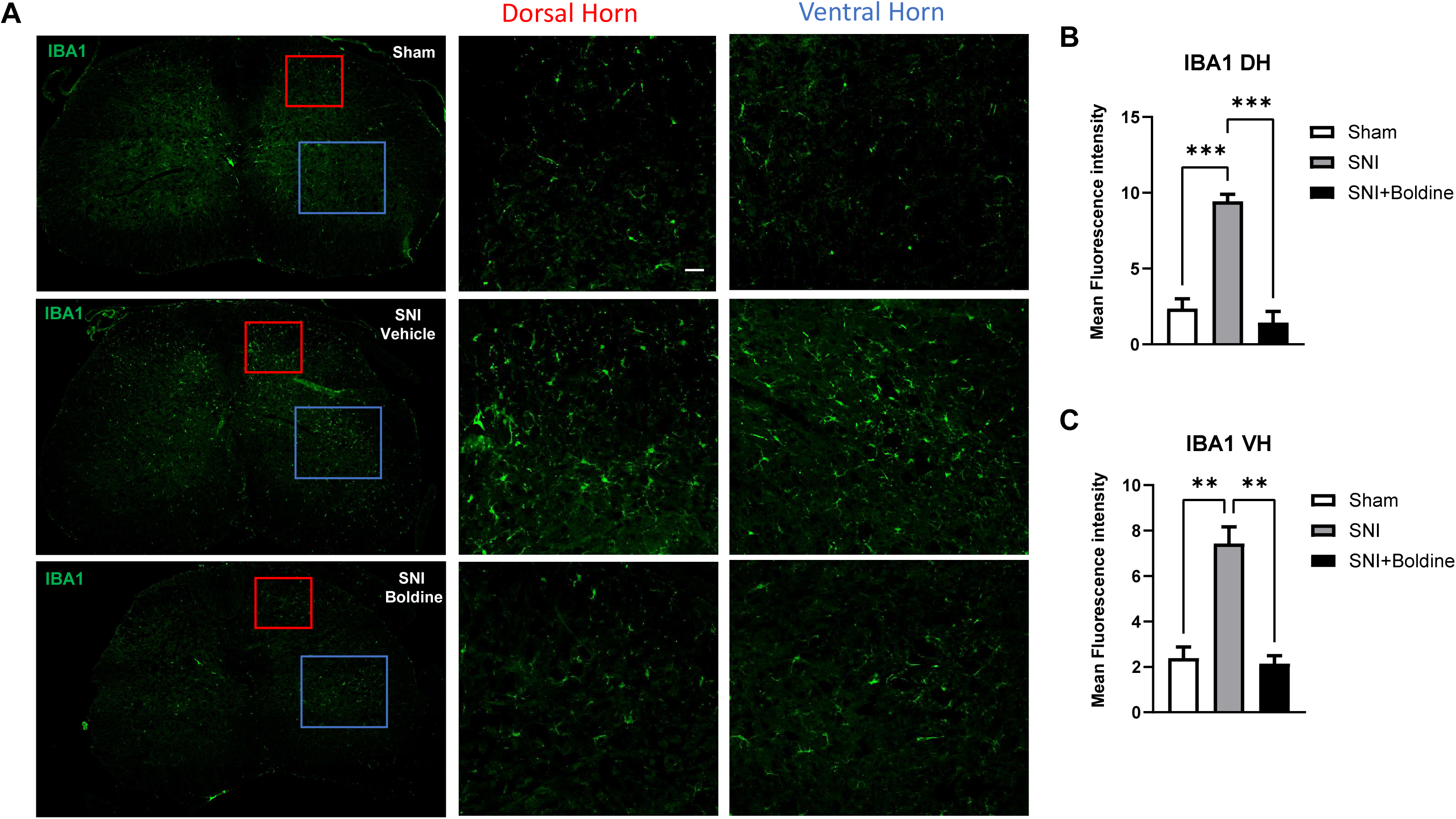
Boldine decreases levels of activated microglia in the lumbar spinal cord after SNI. Sixteen micrometer thick transverse lumbar spinal cord sections were immuno-stained with microglia marker IBA1 to evaluate the effect of SNI and boldine treatment on glia activation. Images were acquired using the ZEISS LSM 700 confocal imaging system with 20 × objective. (A) Representative images of the lumbar spinal cord. Red box: ipsilateral dorsal horn; Blue box: ipsilateral ventral horn. Scale bar: 10 µm. Quantification of IBA1 expression at the (B) ipsilateral DH and (C) VH. Data represent Mean ± SEM, N = 3 mice per group. ** p < 0.01, *** p < 0.001 by one-way ANOVA with Tukey post-test.

### Boldine inhibits the elevation of neuroinflammation markers in the spinal cord

Glia mediators, such as IL-1β and tumor necrosis factor-α (TNF-α), are key factors in the glial control of pain. It has been reported that overexpression of IL-1β in the injured sciatic nerve following nerve injury may be a common mechanism underlying the generation of neuropathic pain [21]. C-C motif chemokine ligand 2 (CCL2), also known as monocyte chemotactic protein 1 (MCP-1), and its receptor CCR2 may play a role in neuroinflammation by participating in neuron-glial cell interactions [6; 40; 57]. To evaluate the effect of boldine on these inflammatory markers in the spinal cord following SNI, we extracted RNA from lumbar spinal cord collected at 15 dpi and performed quantitative PCR using the standard Taqman method. Significantly elevated levels of TNF-α, IL-1β and CCL2 (Figure 8A-C) mRNA were detected in the lumbar spinal cord following SNI. Oral administration of boldine decreased the levels of TNF-α (Figure 8A) and IL-1β (Figure 8B). We also examined the mRNA levels of the receptors of TNF-α, IL-1β and CCL2, and found no difference (Figure 8D-F).

**Figure 8.**
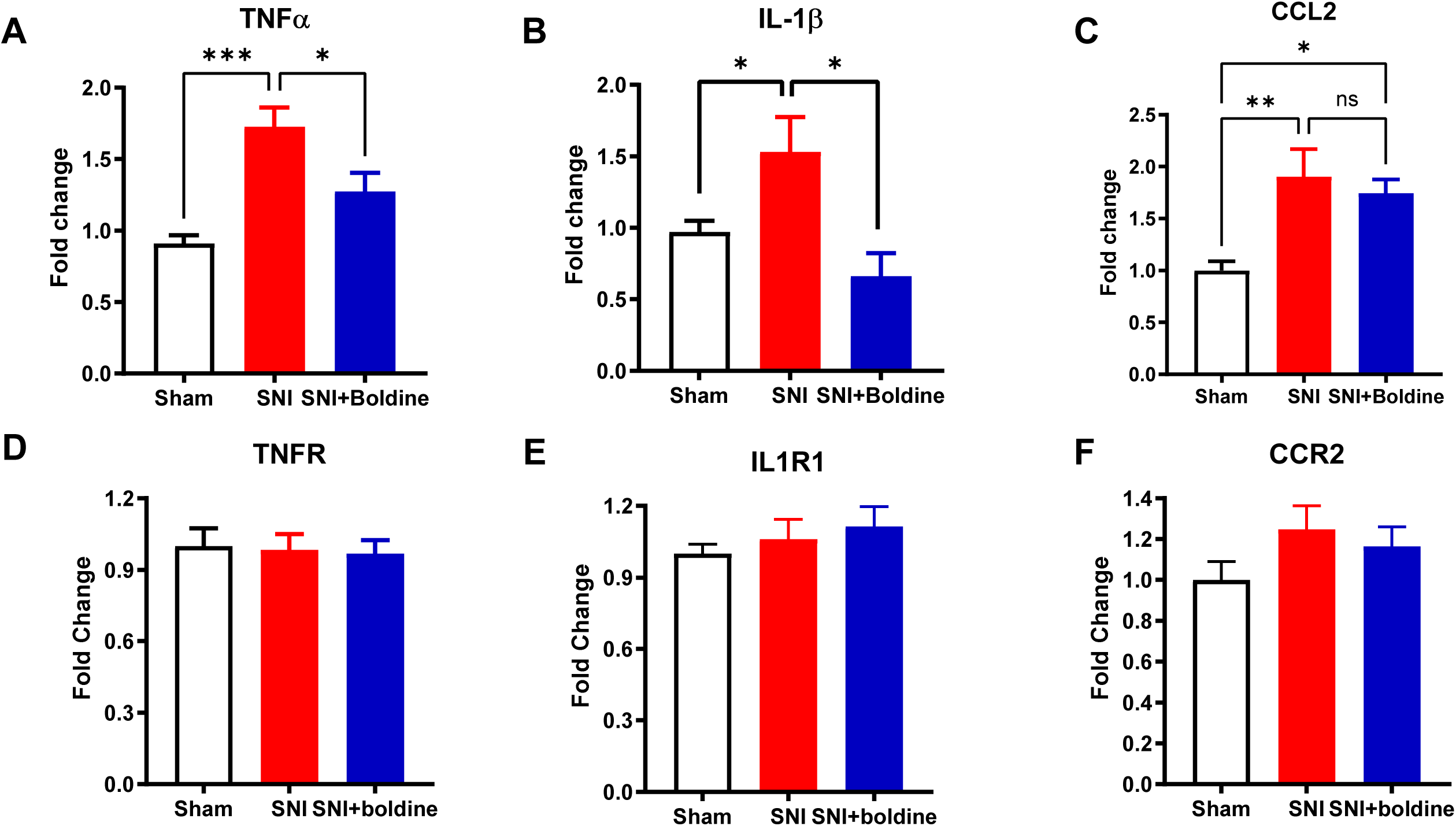
Boldine decreases mRNA levels of inflammation markers in the spinal cord. Total RNA was extracted from lumbar spinal cords collected at 15 dpi and subjected to gene expression analysis by quantitative PCR using standard Taqman methods. Relative levels were calculated using the 2^−ΔΔCt^ method with HPRT as an internal control. (A) TNF-α; (B) IL-1β; (C) CCL2; (D) TNFR; (E) IL1R; (F) CCR2 mRNA levels. Data represent Mean ± SEM, N = 6 per group, * p < 0.05, ** p < 0.01, *** p < 0.001 by unpaired t-test.

## Discussion

The SNI model is a commonly used model in rodents to study neuropathic pain. SNI leads to mechanical allodynia, which develops 3 days after injury and lasts between 4 weeks to 12 months in mice [22]. SNI also leads to thermal hypersensitivity and weight bearing deficits. In this study, we administered boldine via a customized diet to avoid potential stress associated with gavage or injections. Boldine is a bitter alkaloid and there is no added flavor in the diet, one potential risk is that the bitter taste might negatively affect food intake in the boldine treatment group. We monitored food consumption and did not observe any difference between experimental groups, suggesting that the bitter taste does not have any influence on food intake. We found that oral administration of boldine could prevent SNI-induced mechanical allodynia at 3, 7, and 14 dpi. A potential confound is whether boldine could impair locomotor function which could slow or weaken paw withdrawal responses to mechanical stimulus. Several lines of evidence argue against this possibility. First, boldine did not alter the withdraw threshold on the contralateral hind paw; Second, a recent study from our lab showed no effect of boldine on paralyzed muscle weights [47]; Third, the pharmacological effects of boldine have been studied in naive and multiple rodent models of disease without any reported defects in motor function [48]. Boldine prevented SNI-induced thermal hypersensitivity and weight bearing deficit at 7 dpi. We did not observe boldine-mediated prevention of weight bearing deficits at 3 dpi, which might be partially attributed to possible neophobia to the diet (i.e. receiving a lower dose of boldine for the first several days). We also noticed that animals started to recover from thermal or weight bearing pain responses at 14 dpi, which is similar to previous report that thermal hypersensitivity and weight bearing share similar pain recovery profiles, whereas mechanical allodynia persists in a sciatic nerve entrapment injury model [52]. At 14 dpi, the ipsilateral/contralateral hindlimb weight support ratios were similar among experimental groups, while the support area ratios remained significantly lower in injured animals, suggesting a more persistent and slower recovery of weight support area deficits.

Inflammatory responses are of great significance in the development of neuropathic pain. A large body of evidence indicates that inflammatory mediators in local inflamed tissues, peripheral nerves, and spinal cord make important contributions to the initiation and maintenance of chronic pain [27; 29]. In this study, we have observed significantly elevated expression of cytokine/chemokines including IL-12p40, IL-15, CCL5, and M-CSF in the serum after SNI. Subcutaneous injection of IL-12 leads to mechanical hyperalgesia in rats [58]. IL-15 has been linked with nerve injury-induced neuropathic pain in a rodent CCI models [18] as well as rheumatoid arthritis- and osteoarthritis-induced joint pain in humans [61]. Intradermal injection of CCL5 has been shown to induce pain and administration of Met-CCL5, an antagonist of CCL5 receptor CCR5, attenuates inflammatory and nociceptive responses following peripheral nerve injury [38]. In humans, upregulated CCL5 has been detected in patients with various pain conditions including small fiber neuropathy [32], atypical facial pain and trigeminal neuralgia [35], and discogenic back pain [19; 33]. M-CSF is also an important contributing factor to the development of neuropathic pain [64] and arthritic pain [11]. Our data suggested that boldine administration prevented the SNI-mediated induction of these pain-related inflammatory mediators in the periphery.

Neuroglial cells, including microglia, astrocytes, oligodendrocytes, and ependymal cells, play complex and dynamic roles in pain processing. Their activation and interactions with neurons contribute to the amplification and maintenance of pain signals, as well as the development of chronic pain conditions [26]. Our immunofluorescence data confirmed an increased number of reactive astrocytes and activated microglia in the ipsilateral spinal cord. Boldine treatment significantly prevented the injury-induced glia activation. Notably, we also observed some levels of reactive astrocytes in the contralateral spinal cord sections at 15 dpi (Figure 7A), which could possibly be attributed to the central sensitization process. Recent studies have shown that after nerve injury, toll-like receptors (TLRs) are involved in the generation of neuropathic pain. For example, TLR2 and TLR4 modulate glial activation in the spinal cord and contribute to neuropathic pain development [37; 54]. TLRs are responsible for the stimulation of astrocytes and microglia, which will induce proinflammatory mediators and cytokines, therefore leading to the generation and maintenance of neuropathic pain. The majority of TLRs can signal primarily through the myeloid differentiation 88 (MyD88) pathway, and then trigger the activation of ERK/JNK/NFκB signaling. Nod-like receptors (NLRs) are cytoplasmic sensors that oligomerize to form the inflammasome, a multi-subunit complex that processes the maturation of pro-IL-1β and pro-IL-18 via caspase-1 [23]. A well-characterized outcome of TLR and inflammasome interaction is the secretion of IL-1β and IL-18 [49]. In this study, we report that boldine prevented the LPS-induced overexpression of multiple inflammatory cytokines and chemokines including CCL2, IL-1β, IL-6, and IL-18 in BV-2 microglial cells. We have also found decreased expression of toll-like receptors (e.g. TLR2, TLR3), and NLRP3 after boldine treatment, suggesting that boldine might prevent neuroinflammation through inhibiting the TLR/NLRP3 inflammasome signaling. Significant decrease of TNF-α and IL-1β mRNA in the lumbar spinal cord was detected in boldine-treated animals at 15 dpi. The boldine-mediated changes at the protein level and the direct target of boldine still remain to be characterized.

In summary, we report for the first time that oral administration of boldine, a naturally-occurring alkaloid, attenuated neuropathic pain after spared nerve injury, suggesting that boldine might be considered as a novel therapeutic agent for the treatment of neuropathic pain. Future studies will explore how the boldine and its derivatives differentially influence pain perception in the central nervous system and peripheral nervous system across sexes and extend our research to other pain conditions, including arthritic pain, back pain, and diabetic pain.

## Author Contributions

Experiment design: WZ, CPC; experiment execution and data collection: JP, WZ, CAT; data analysis: WZ, JP, CC, CAT; manuscript writing and editing: WZ, CPC, JCS, YD.

## Acknowledgment

The authors would like to thank Dr. Miguel Gama-Sosa for helpful discussions related to technical aspects of the studies reported.

## Abbreviations

ANOVA: Analysis of variance
BW: body weight
CCI: Chronic constriction injury
CCL: C-C Motif Chemokine Ligand
CCR: C-C Motif Chemokine receptor
CXCL: C-X-C motif chemokine ligand
DH: dorsal horn
DMEM: Dulbecco’s Modified Eagle Medium
Dpi: Days post injury
DWB: Dynamic Weight Bearing
ELISA: enzyme-linked immunosorbent assay
FBS: fetal bovine serum
GFAP: Glial fibrillary acidic protein
HPRT: Hypoxanthine phosphoribosyltransferase 1
IL: Interleukin
IL1R: IL-1 receptor
LPS: Lipopolysaccharides
M-CSF: macrophage-colony stimulating factor
NFκB: nuclear factor-kappa B
NLRP3: Nod-like receptor pyrin containing 3
PBS: Phosphate-buffered saline
PFA: paraformaldehyde
SEM: standard error of the mean
SNI: Spared nerve injury
TNF-α: Tumor Necrosis Factor Alpha
TNFR: Tumor Necrosis Factor Receptor
TLR: Toll-like receptor
VH: ventral horn

