## Supplementary files for "Oral Administration of Boldine Reduces Spare Nerve Injury-Induced Neuropathic Pain"

**Supplemental Table 1. List of Taqman probes for qPCR.**

| Gene | Probe |
|---|---|
| CCL2 | Mm00441242_m1 |
| CCL3 | Mm00441259_g1 |
| CCL5 | Mm01302427_m1 |
| CCR2 | Mm99999051_gH |
| CD11b | Mm00434455_m1 |
| HPRT | Mm03024075_m1 |
| IL-1β | Mm00434228_m1 |
| IL1R | Mm00434237_m1 |
| IL-6 | Mm00446190_m1 |
| IL-18 | Mm00434226_m1 |
| NLRP3 | Mm00840904_m1 |
| TLR2 | Mm01213946_g1 |
| TLR3 | Mm01207404_m1 |
| TNF-α | Mm00443258_m1 |
| TNFR | Mm00441883_g1 |

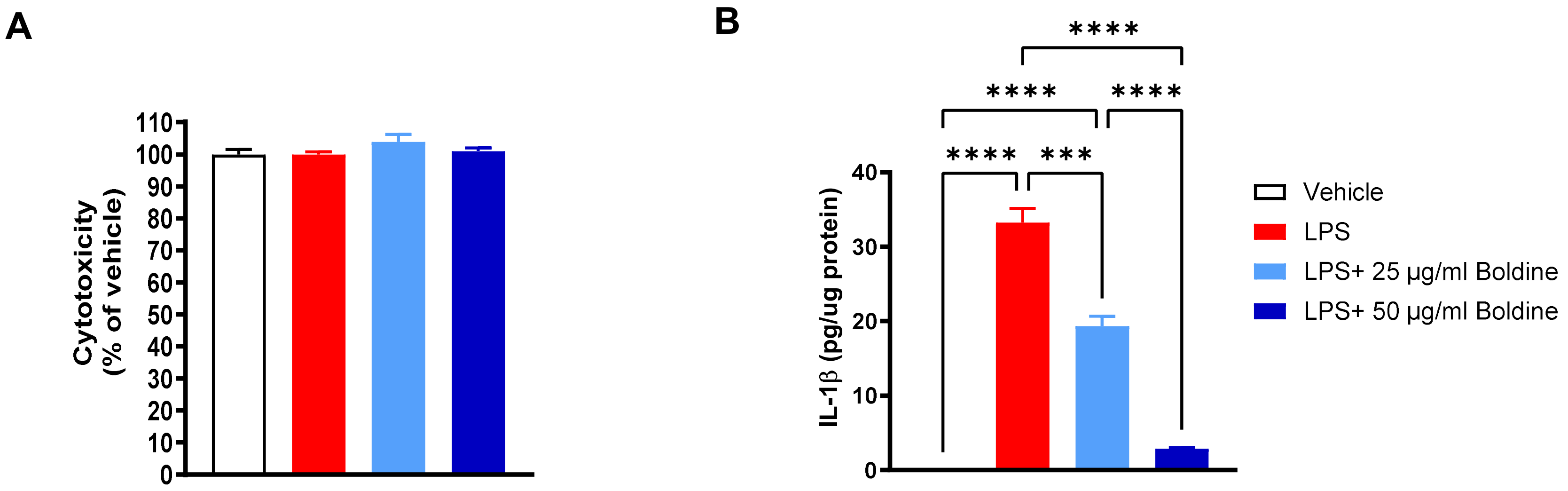


**Supplemental Figure 1. Effect of boldine on cytotoxicity and IL-1β protein expression.** BV-2 microglia cells were treated with 100 ng/mL lipopolysaccharides (LPS) with or without boldine (25 μg/mL or 50 μg/mL) for 6 hours. After treatment, cell culture medium was collected, and cytotoxicity was assessed using the CytoTox96 Non-Radioactive Cytotoxicity Assay (Promega). Cells were harvested, lysed in RIPA buffer and IL-1β protein levels were assessed using the Mouse IL-1 beta/IL-1F2 Quantikine ELISA Kit (R&D Systems). (A) Boldine treatment did not lead to any change in cell viability; (B) Boldine significantly reduced IL-1β protein levels in the cell lysates. Data represent Mean ± SEM, N = 3 per condition, *** $p < 0.001$, **** $p < 0.0001$ by one-way ANOVA followed by Tukey post-tests.

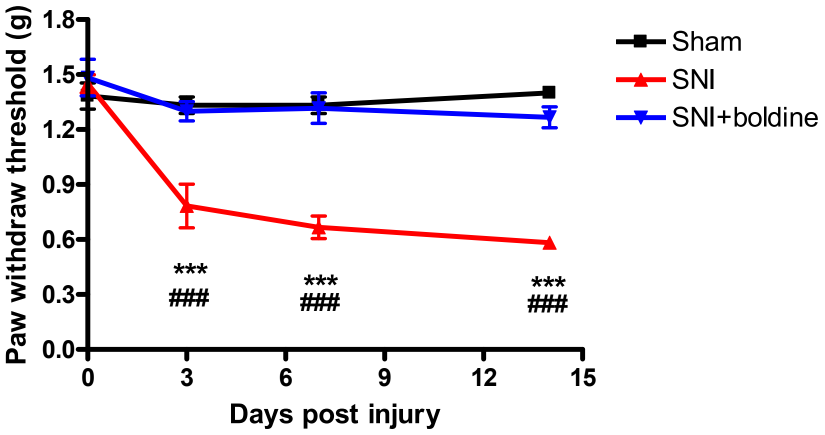


**Supplemental Figure 2. Boldine reduces SNI-induced mechanical hypersensitivity in male mice.** Ten weeks old male C57BL6 mice were subjected to SNI subjected to spared nerve injury (SNI). Boldine was administered right after injury via a customized diet (Research Diets) to reach a dose of 50 mg/kg/day. A control diet was provided to sham or SNI group. Data represent mean±SEM, N = 8-10 per group, *** $p < 0.001$ (SNI vs. sham), ### $p < 0.001$ (SNI+boldine vs. SNI) by two-way ANOVA with Tukey post-tests.
